# A novel density-based neural mass model for simulating neuronal network dynamics with conductance-based synapses and membrane current adaptation

**DOI:** 10.1101/2020.10.09.334144

**Authors:** Chih-Hsu Huang, Chou-Ching K. Lin

**Affiliations:** Department of Neurology, National Cheng Kung University Hospital, College of Medicine, National Cheng Kung University, Tainan, Taiwan

## Abstract

Nowadays, building low-dimensional mean-field models of neuronal populations is still a critical issue in the computational neuroscience community, because their derivation is difficult for realistic networks of neurons with conductance-based interactions and spike-frequency adaptation that generate nonlinear properties of neurons. Here, based on a colored-noise population density method, we derived a novel neural mass model, termed density-based neural mass model (dNMM), as the mean-field description of network dynamics of adaptive exponential integrate-and-fire neurons. Our results showed that the dNMM was capable of correctly estimating firing rate responses under both steady- and dynamic-input conditions. Finally, it was also able to quantitatively describe the effect of spike-frequency adaptation on the generation of asynchronous irregular activity of excitatory-inhibitory cortical networks. We conclude that in terms of its biological reality and calculation efficiency, the dNMM is a suitable candidate to build very large-scale network models involving multiple brain areas.

## Introduction

Instead of exploring spiking activity of individual neurons, sometimes, scientists are more interested in the summed activity of a neuronal population, composed of thousands or ten thousands of neurons, in response to external stimuli for several reasons. One reason comes from the accepted concept that the collective activity of interconnected neurons within multiple brain areas is considered to account for brain functions, such as sensory, motor, and cognitive functions (***Gerstner et al., 2014; Deco et al., 2015***). Another reason is that functional measurements of brain states, for examples, recorded from functional magnetic resonance imaging and electroencephalography, reflect the collective activity of thousands of neurons (***Deco et al., 2008; Breakspear, 2017***). In addition, investigating network dynamics of neuronal populations at mesoscopic scales bridges the microscopic states of individual neurons to the macroscopic global brain states (***Goldman et al., 2019***). Building computational models of such neuronal populations thus has been one of the major goals in computational neuroscience.

The computational modeling of neuronal populations can be achieved by means of bottom-up theoretical modeling approaches, ranging from direct numerical simulations of extremely detailed spiking neurons (***Izhikevich and Edelman, 2008; Markram et al., 2015***) to more coarse-grained rate models (***Dayan and Abbott, 2001***), or neural mass models (***David and Friston, 2003; Moran et al., 2013***). The direct numerical simulation of spiking neurons encompasses essential details of individual neuronal and synaptic dynamics. Such a modeling approach thus requires huge computational resources, and it is really difficult to analyze the dynamic behaviors of such a large system of neural networks. Therefore, mathematical models directly describing the collective activity of neuronal populations emerge under the mean-field assumptions to reduce the computational demands (***Deco et al., 2008; Breakspear, 2017***).

The neural mass model (NMM) describes the mean activity of a neuronal population, which is derived by assuming that in the presence of strong coherence among neurons, the collective activity is sufficiently close to the mean so that the variance can be discarded (***Breakspear, 2017***). Those mean quantities typically include population-average firing rates, mean membrane potentials and mean synaptic state variables, whose dynamic behaviors are described by a single or small number of ordinary differential equations and, in general, a sigmoid rate function transferring the mean membrane potential into the population firing rate. The history of the NMM can be traced back to 70’s (***Lopes da Silva et al., 1974; Zetterberg et al., 1978***), and it has been seen in many examples of simulating various EEG rhythms during both the wakefulness and sleep (***Cona et al., 2014***), the generation of resting-state functional magnetic resonance imaging data (***Deco et al., 2013***), or pathological activity of neurological diseases (***Suffczynski et al., 2004; Chiang et al., 2011; Bhattacharya et al., 2011***). Moreover, it is now incorporated in the Virtual Brain Project that aims to deliver the first open simulation of the human brain based on the large-scale connectivity of neurons (***Sanz-Leon et al., 2015***).

However, an opposing opinion criticizes that the NMM is essentially phenomenological, possibly only reproducing a part of the rich repertoire of dynamic responses seen in real neuronal tissues (***Coombes and Byrne, 2019***). Furthermore, despite of its success, some disadvantages of NMMs also have been demonstrated recently. For examples, NMM lacks systematic link to microscopic models at the level of spiking neurons (***Schwalger et al., 2017***). Second, it fails to correctly reproduce the firing rate responses of a population of neurons when the input changes rapidly, although it has good accuracy of predicting steady-state responses (***Deco et al., 2008; Gerstner et al., 2014***). Third, fluctuations of activity variables in NMMs are absent. Forth, the use of a sigmoid rate function for estimating population activity is not derived from a biophysically detailed description of spiking neurons (***Byrne et al., 2020***). Accordingly, a new version of NMMs is mandatory.

To be biologically realistic, we focused on the development of a novel NMM for the modeling of a population of adaptive exponential integrate-and-fire (aEIF) neurons (***Brette and Gerstner, 2005***), in which two critical neuronal features, i.e., adaptive behaviors of firing rate responses and nonlinear voltage-dependent synaptic interactions, were included. Experiments showed that slowly adapting spike-frequency responses, a pattern called regular spiking (RS), were generally observed in virtually all excitatory neurons in cerebral cortex, while inhibitory neurons often fired at higher frequencies with no adaptation, a pattern called fast spiking (FS) (***Connors and Gutnick, 1990***). Nonlinearity of voltage-dependent synaptic interactions also has been displayed to play a key role in network responses to external stimuli (***Paulus and Rothwell, 2016; Zerlaut and Destexhe, 2017; di Volo et al., 2019***). Such voltage-dependent synaptic interactions were modeled in a conductance-based manner in our study.

Our work was based on the so-called colored-noise population density method (csPDM), proposed by ***Huang and Lin (2018)***, where the authors yielded a Fokker-Planck formalism to govern the temporal evolution of the probabilistic distribution of membrane potential state variables across a population of exponential integrate-and-fire (EIF) neurons (***Fourcaud-Trocmé et al., 2003***), together with conductance-based synaptic interactions, and then, to evaluate the dynamic population firing rate behavior in response to external inputs. Here, we expanded it to the case where adaptation was included. It is well known that the solution of the Fokker-Planck formalism was shaped by a Gaussian density function which could be completely described by the associated statistical mean and variance (***Risken, 1996***). In other words, the distribution of membrane potential state variables in the csPDM could also be well characterized by the corresponding mean and variance values. As a result, by deriving the dynamic equations for evaluating the temporal evolution of the mean and variance from the Fokker-Planck formalism, the csPDM could be transformed into a type of NMMs together with a semi-analytic rate function. In this study, we illustrated the transformation from the csPDM into the NMM based on this idea. Owing to its inspiration from the csPDM, we named this novel NMM the density-based NMM (dNMM).

In addition to treating the drawbacks of conventional NMMs, our dNMM was designed also to tackle the problem of computational inefficiency of population density methods (***Tranchina, 2009***). When building a large-scale network model consisting of multiple neuronal populations, the population density method lost its advantage as a time-saving alternative to direct numerical simulations of detailed spiking neuron networks, even though certain dimension-reduction algorithms were employed for each single neuronal population (***Apfaltrer et al., 2006***). Under this circumstance, however, our dNMM remained highly efficient in calculation because the dynamic equations tracing state variables were in the forms of ordinary differential equations, rather than partial differential equations. As such, it could be expected that the dNMM reasonably fit in the whole brain model for simulating the resting-state behaviors seen in BOLD signals (***Deco et al., 2013***).

In the current study, the dNMM was demonstrated to have excellent accuracy to reproduce steady or transient firing rate responses of a single aEIF neuronal population with different strengths of spike-frequency adaptation. In particular, we utilized the dNMM to investigate the network dynamics of an excitatory-inhibitory network, namely, the standard minimal circuitry for cortical neuronal networks (***Okun and Lampl, 2008; Avermann et al., 2012***). The modeled dynamics focused on the regime of asynchronous irregular (AI) activity that was experimentally observed in mammalian cortex under awake states (***Matsumura et al., 1988; Brunel, 2000; Steriade et al., 2001; Destexhe et al., 2003; Lee et al., 2006; El Boustani et al., 2007***). We explored whether the spike-frequency adaptation affected the extent of AI regimes in a parameter space spanned by two control parameters, that is, the amplitude of external input and the relative strength of inhibitory synaptic interactions. We showed that such adaptation enlarged the AI regime in the parameter space, suggesting that the spike-frequency adaptation in the cortical neurons possibly played a critical role in maintaining the AI activity under the fluctuation of external inputs and neuronal properties.

## Results

### Dynamic equations of density-based neural mass models

We were interested in the firing rate dynamics of a large homogeneous population of *N* uncoupled aEIF neurons in response to external excitatory and inhibitory Poissonian spike trains. The synaptic interaction was handled by using conductance-based methods with the first-order kinetics of synaptic conductances. In this case, the state of each aEIF neuron was described by four state variables, i.e., the membrane potential, *V*(*t*), adaptation current, *w*(*t*), excitatory and inhibitory synaptic conductances, *g_e_*(*t*) and *g_i_*(*t*), which temporally evolved in response to overall presynaptic excitatory and inhibitory spike inputs. Assuming that the mean rates of the excitatory and inhibitory Poisson spike inputs were *v_e_*(*t*) and *v_i_*(*t*), respectively, the dynamic equations of the dNMM were:

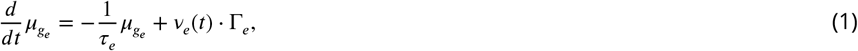

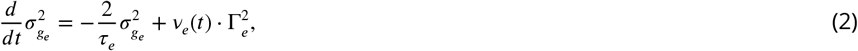

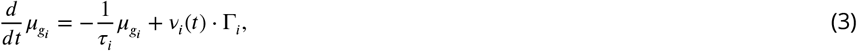

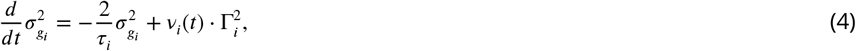

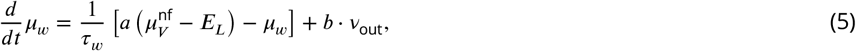

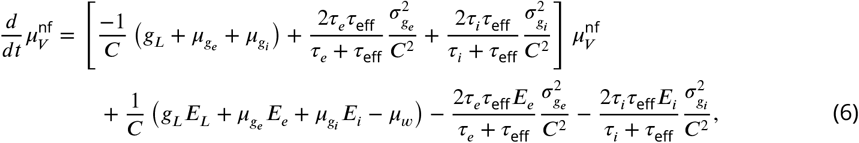

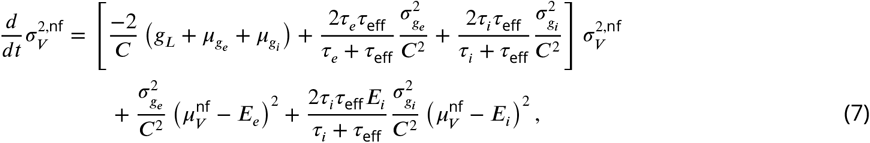

where *μ*_*_ denoted the associated means of state variables, and 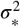 the variances; *C* was the membrane capacitance, *g_L_* the leak (resting) conductance, *E_L_* the leak reversal potential. *τ*_eff_ was the effective membrane time constant, *τ_w_* the time constant of the adaptation current, *τ_e_* the time constant of the excitatory synaptic dynamics and *τ_i_* the inhibitory one. Here, Γ_*e*_ and Γ_*i*_ stood for the synaptic strengths, which were, respectively, the jump sizes of the excitatory and inhibitory synaptic conductances caused by one presynaptic spike under the assumption of 1-exponential synaptic kinetics (***Roth and van Rossum, 2009***). The mean adaptation current *μ_w_* was controlled both by the subthreshold adaptation parameter *a* and the spike-frequency adaptation parameter *b. v*_out_ was the so-called population-average firing rate of the neuronal population. Notice that although the neuronal model was conductance-based, *v*_out_ was computed by a current mapping algorithm, which was given by:

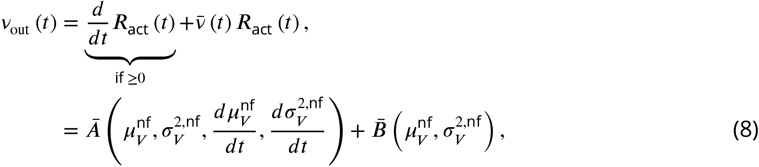

where

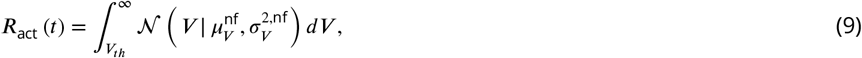

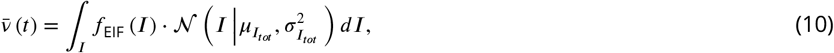

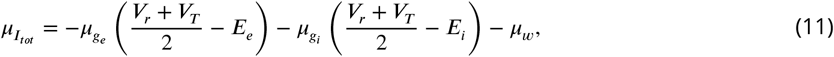

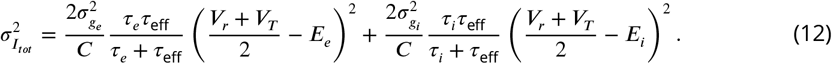

Here, *Ā*(*) accounted for transient changes of the firing rate in response to the rapidly changing input, while 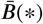 expressed the steady firing rate. *R*_act_(*t*) denoted the proportion of activated neurons whose membrane potentials exceeded a certain threshold voltage *V_th_* within the neuronal population at the present time *t*. 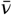 was the estimate of the mean firing rate of those activated neurons. 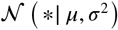 meant the Gaussian density function with the mean *μ* and variance *σ*^2^. *f*_EIF_(*I*) was the current-rate relationship of a single EIF neuron that mapped the current input into its mean output firing rate, which, in this study, was obtained numerically before running simulations. *μ_I_tot__* denoted the total mean effective input current received by a neuron, and 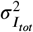 meant the variance of the effective input current. It should be noted that the superscript “nf” meant that the mean and variance of membrane potentials were estimated under the removal of spike-firing mechanisms of individual aEIF neuronal dynamics. In short, the dynamic equations of the dNMM included seven ordinary differential equations and a rate estimate function for evaluating the population-average output firing rate, together with a few auxiliary variables. All descriptions and derivations of the state variables and dynamic equations above are presented in the section of Methods and Materials.

### Gaussian distributions of membrane potential state variables

A core assumption of the dNMM was the Gaussian distribution of membrane potential state variables across the neuronal population. We attempted to explore whether such an assumption was valid with respect to different synaptic strengths and input rates. Before this, the premise of the csPDM, i.e., the diffusion approximations of synaptic dynamics (***Huang and Lin, 2018***), were first checked. Such an approximating technique was often used in the computational studies of network dynamics for considering the fluctuations of synaptic interactions (***Fourcaud and Brunel, 2002; Richardson, 2004; Linaro et al., 2011; Augustin et al., 2013; Rosenbaum, 2016; Augustin et al., 2017***). In our cases, the steady distributions of synaptic conductances were postulated to be shaped by Gaussian density functions under diffusive limits, i.e., the input rate *v_e,i_* ≫ 1 and the synaptic strength Γ_*e,i*_ ≪ 1. First, we checked the reasonable physiological ranges of synaptic strengths. As shown by the blue line in ***Figure 1A***, the peak voltage jump Γ_*V*_ caused by individual excitatory presynaptic spikes revealed a linear relationship with the amplitude of the synaptic strength Γ_*e*_. In particular, Γ_*V*_ = 0.5 mV (a typical value for a large network of 10,000 to 100,000 neurons (***El Boustani et al., 2007; Brémaud et al., 2007***)) when Γ_*e*_ = 0.0025 mS. So, we explored the distributions of excitatory synaptic conductances with three different synaptic strengths of 0.0025, 0.005 and 0.01 mS which gave rise to peak voltage jumps of 0.5, 1 and 2 mV, respectively. The input rate for each case was just at the half of *v_c′_*, which was the minimum input rate of an excitatory Poisson spike train whose mean input allowed the aEIF neuron to repeatedly emit spikes, given by

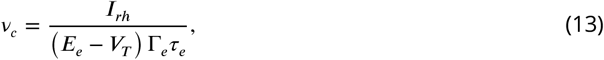

where *I_rh_* was defined as the rheobase current that was the minimum constant current required to elicit a spike (see the Eq. (7) in ***Touboul and Brette (2008)***).

**Figure 1.**
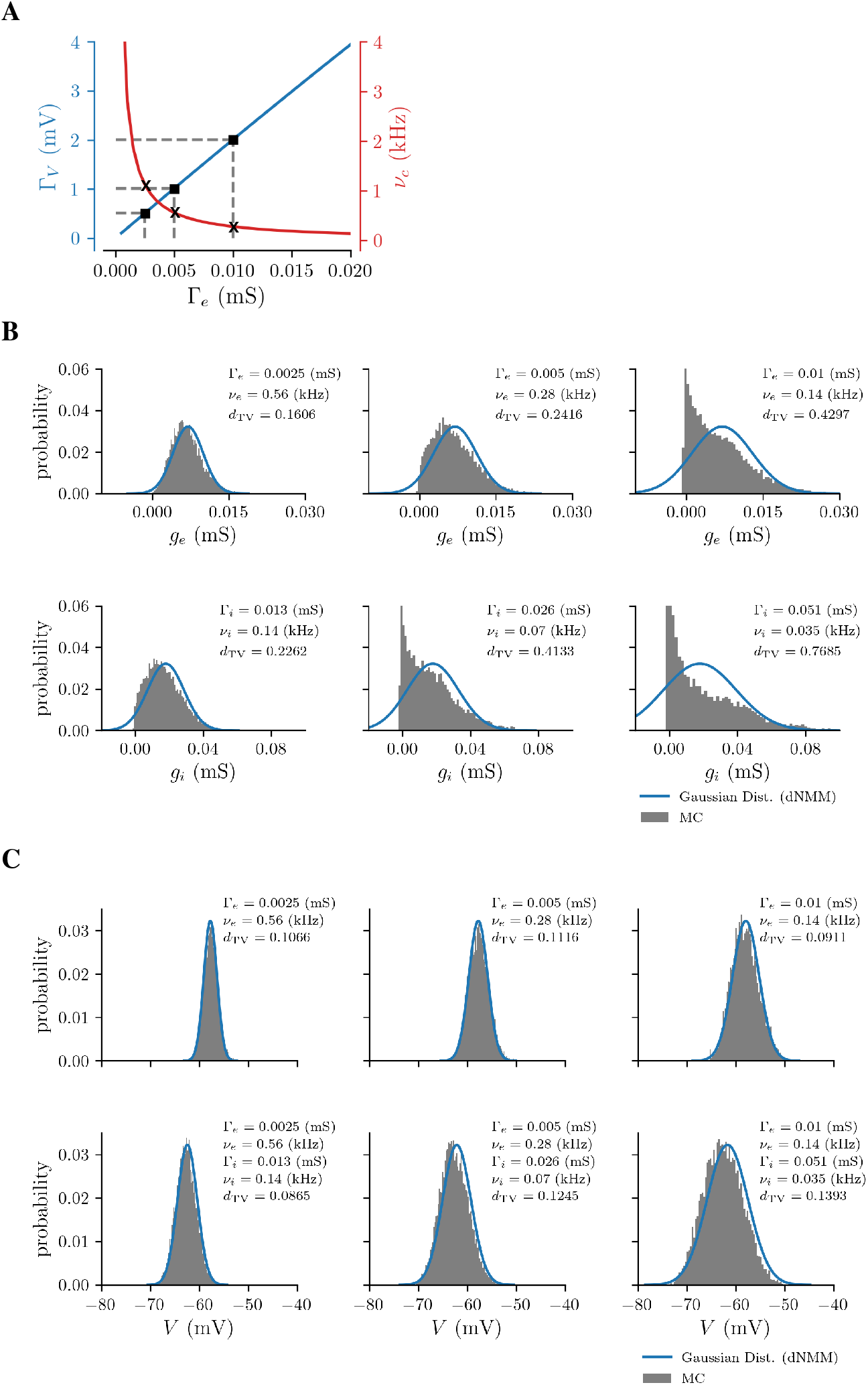
Validity of diffusion approximations of synaptic dynamics. (**A**) Relationships of excitatory synaptic strengths Γ_*e*_ with the peak voltage jump sizes Γ_*V*_ caused by individual presynaptic spikes (left axis, the blue line) and the minimum input rates *v_c_* required to emit spikes (right axis, the red line). The black squares indicate the Γ_*e*_ chosen for validating diffusion approximations, and the corresponding *v_c_* are indicated by x notations. (**B**) Comparisons of steady distributions of postsynaptic conductances with Monte-Carlo (MC) simulations, i.e., direct numerical simulations of synaptic dynamics. The upper row is for excitatory conductances and the bottom row for inhibitory conductances. (**C**) Comparisons of steady distributions of membrane potentials with Monte-Carlo (MC) simulations of aEIF neuron networks. The upper row shows the distributions of membrane potentials of aEIF neurons only receiving excitatory input, while the bottom row shows the associated distributions where aEIF neurons receive excitatory and inhibitory inputs. The corresponding steady distributions of either excitatory or inhibitory postsynaptic conductances are shown in **B**.

The upper row in ***Figure 1B*** displays steady distributions of excitatory synaptic conductances from Monte-Carlo (MC) simulations, i.e., direct numerical simulations of synaptic dynamics as ground truths (shaded areas), and Gaussian density functions from diffusion approximations whose means and variances were given by equilibrium points of Eqs. (1) and (2) (blue lines). As expected, the diffusion approximation was in good agreement with the MC simulation only in the case of Γ_*e*_ = 0.0025 mS and *v_e_* = 0.56 kHz, that is, under the condition of small synaptic strengths and large input rates. To highlight this, we quantitatively characterized the effectiveness of diffusion approximations by using the total variation of histogram probabilities, given by

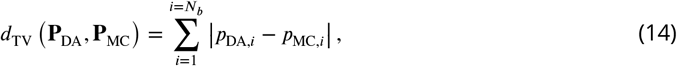

where *N*_b_ was the number of bins, **P**_DA_ meant a histogram probability vector consisting of *N_b_* elements, i.e., *p*_DA,*i*_, where *i* = 1,2,…, *N_b_*, which was the histogram probability within the *i*-th bin obtained from the diffusion approximation; similarly, **P**_MC_ for MC simulations. We set *N_n_* = 100 bins ranging from *μ_g_e__* – 4*σ_g_e__* to *μ_g_e__* + 4*σ_g_e__*. In the case of Γ_*e*_ = 0.0025 mS and *v_e_* = 0.56 kHz, *d*_TV_ was 0.1606. Through this observation, the diffusion approximation would reveal good estimations of synaptic conductances if *d*_TV_ was below 0.15. For the other two cases of Γ_*e*_ = 0.005 and 0.01 mS, the conductance distributions from MC simulations were markedly different from Gaussian functions, whose *d*_TV_ were 0.2416 and 0.4297, respectively.

We also checked the inhibitory synaptic dynamics. The inhibitory synaptic strength was determined according to the relative strength of effective inhibition, which was given by (***Kumar et al., 2008***):

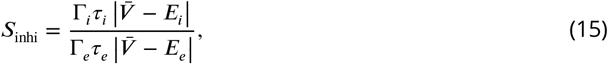

where 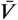 was set as (*V_r_* + *V_T_*)/2 (*V_T_* was the spiking threshold voltage and *V_r_* was the resetting voltage of the aEIF neuron). We let *S*_inhi_ = 4 in order to balance the excitatory and inhibitory recurrent current inputs because the ratio between the excitatory and inhitory recurrent connections was four (***Brunel, 2000; Kumar et al., 2008***). Accordingly, Γ_*i*_ were 0.013, 0.026 and 0.051 mS, respectively, for Γ_*e*_ = 0.0025, 0.005, 0.01 mS. As shown in the bottom row in ***Figure 1B***, we found that *d*_TV_ > 0.2 for all cases because of high synaptic jump sizes and small input rates (*v_i_* was a quarter of *v_e_* in all cases). In other words, the inhibitory synaptic dynamics could not be characterized by diffusionapproximations very well for these cases.

Finally, we assessed the distributions of membrane potentials under those synaptic conditions, where only excitatory inputs or combinations of excitatory and inhibitory inputs were delivered to the neurons. The interesting finding was that, even though, for some cases, the real steady distributions of synaptic conductances did not match Gaussian density functions, the steady distributions of membrane potentials *V* could be correctly captured by Gaussian distributions whose means and variances were estimated through Eqs. (6) and (7) for all cases. The results showed that *d*_TV_ was smaller than 0.15 in all cases (see ***Figure 1C***). It was summarized that, when the synaptic strengths Γ_*e/i*_ were within physiological ranges, the poor performance of diffusion approximations of synaptic conductances might not result in the violation of Gaussian distributions of membrane potential state variables of neurons. Consequently, the assumption of Gaussian density functions in Eqs. (9) and (10) seemed reasonable.

### Mismatch of sigmoid rate functions on estimating population firing rates

The main discrepancy between NMM and dNMM was the different rate estimate functions used for determining population firing rates. The conventional NMM employed monotonic sigmoidal functions of mean membrane potentials to estimate population firing rates. That idea was ubiquitous throughout the modeling studies with NMMs, but the use of sigmoid rate functions did not come from a biophysically detailed descriptions of spiking neurons (***Byrne et al., 2020***). Furthermore, as shown in ***Figure 2A***, where steady population firing rates obtained from MC simulations given by a series of combinations of different excitatory and inhibitory Poissonian input rates are displayed, the population firing rate did not only monotonically increase with the increased mean membrane potentials, but also depended on the standard deviations of membrane potentials across the population. So, the monotonic sigmoid rate function of only mean membrane potentials was insufficient to cover the whole scenario. On the other hand, the dNMM included the estimation of the standard deviation of membrane potentials to determine the firing rates, using Eq. (8). The upper plot in ***Figure 2B*** highlights the difference between the dNMM and NMM in the accuracy of estimating population firing rates, and their estimating errors by comparisons with those from MC simulations are shown in the bottom plot. As seen, the firing rates could be quantitatively correctly estimated by the dNMM. Maximum estimating error of up to ≈ 15 Hz occurred in the regimes where the mean membrane potential 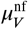 was near the spiking threshold voltage ≈ −50 mV or at the extremely activated region, that is, ≈ −10 mV. All results indicated that the sigmoid rate function, which had a poor estimating accuracy in our cases, was inadequate as an estimate function for determining firing rates. This was due to the fact that, as mentioned-above, it was not derived based on the biophysical understandings of spiking neurons.

**Figure 2.**
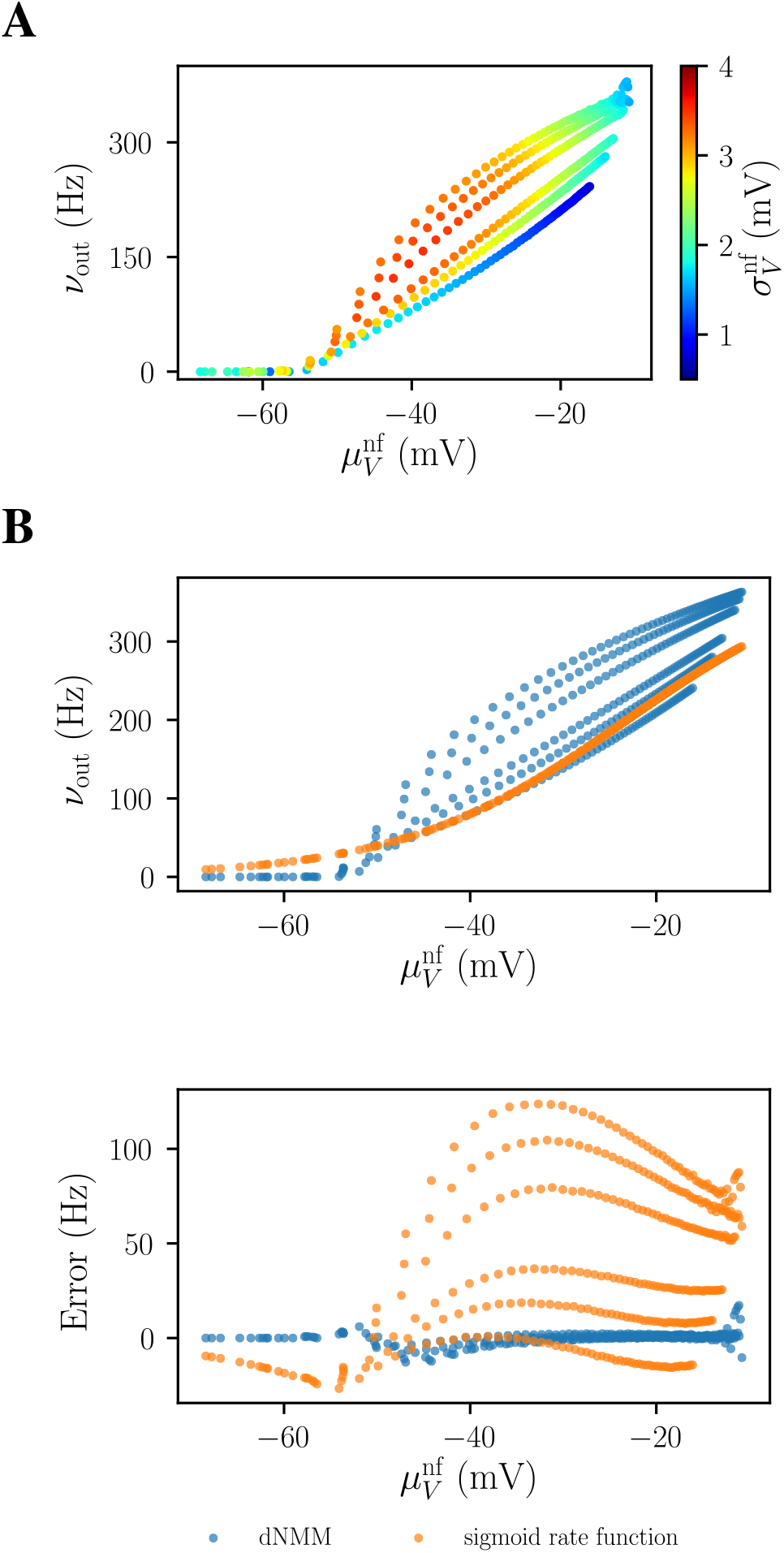
Comparison of dNMMs with conventional NMMs in estimating population firing rates. (**A**) Nonmonotonic relationship of population firing rates with mean membrane potentials. The population firing rates are estimated by MC simulations, and different circles indicate the estimated firing rates with respect to different sets of excitatory and inhibitory Poissonian input rates. The colors of circles map the values of the standard deviations of membrane potentials across the population. *v*_out_: population firing rates; 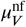: mean membrane potentials; 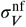: standard deviations of membrane potentials. (**B**) Upper plot: steady population firing rates obtained from the dNMM (blue circles) and conventional NMM (orange circles) given by the same input rate in **A**. Lower plot: the corresponding estimating errors of the dNMM (blue circles) and NMM (orange circles) by comparisons with those from MC simulations. The sigmoid rate function was 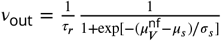, whose parameters were determined by an optimizing procedure (least squares methods) with data from MC simulations given only by excitatory input. Under this circumstance, *μ_s_* = −23.165 mV and *σ_s_* = 12.163 mV.

### Performance of simulating steady-state firing rate responses

Here, and in the next section, we further checked the accuracy of Eq. (8) in estimating population firing rates of neuronal populations with different strengths of spike-frequency adaptation. First, we explored steady-state responses, which were mainly dominated by 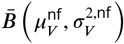. The inhibitory input rates were scanned from 50 to 1000 Hz, and the excitatory input rates were scanned from the fluctuation-driven operating regime, namely, *v_e_*/*v_c_* ≈ 1, in which synaptic-input fluctuations were the primary cause of neuronal firings, to the mean-driven operating one, *v_e_*/*v_c_* ≈ 1, in which neuronal firings were determined primarily by driving inputs (***Cai et al., 2012***). Here, the values of spike-frequency adaptation parameters were set as 0,0.02 and 0.1 for representing adaptation-free, weak adaptation and strong adaptation. As expected, larger spike-frequency adaptation lowered the steady output firing rates under the same input conditions (see the upper row in ***Figure 3***). A counter-intuitive finding was that, under the same excitatory input rate, larger inhibitory input rate led to larger output firing rate. This was due to the increased fluctuations of membrane potentials originating from additional inhibitory input. As compared with those from MC simulations (see the bottom row in ***Figure 3***), it was found that the estimation of firing rates had apparent errors of up-to ≈ 15 Hz when the neurons were driven under fluctuation-driven regimes. But, such errors were smaller for neurons with strong adaptation. The results also showed that under mean-driven operating regimes, the estimated steady-state output firing rates were comparable to those from MC simulations. In summary, the term 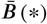 could correctly capture the steady population firing rates under a wide range of input rates, except for the fluctuation-driven operating regimes where the fluctuation magnitude of membrane potentials dominated the accuracy of estimating firing rates.

**Figure 3.**
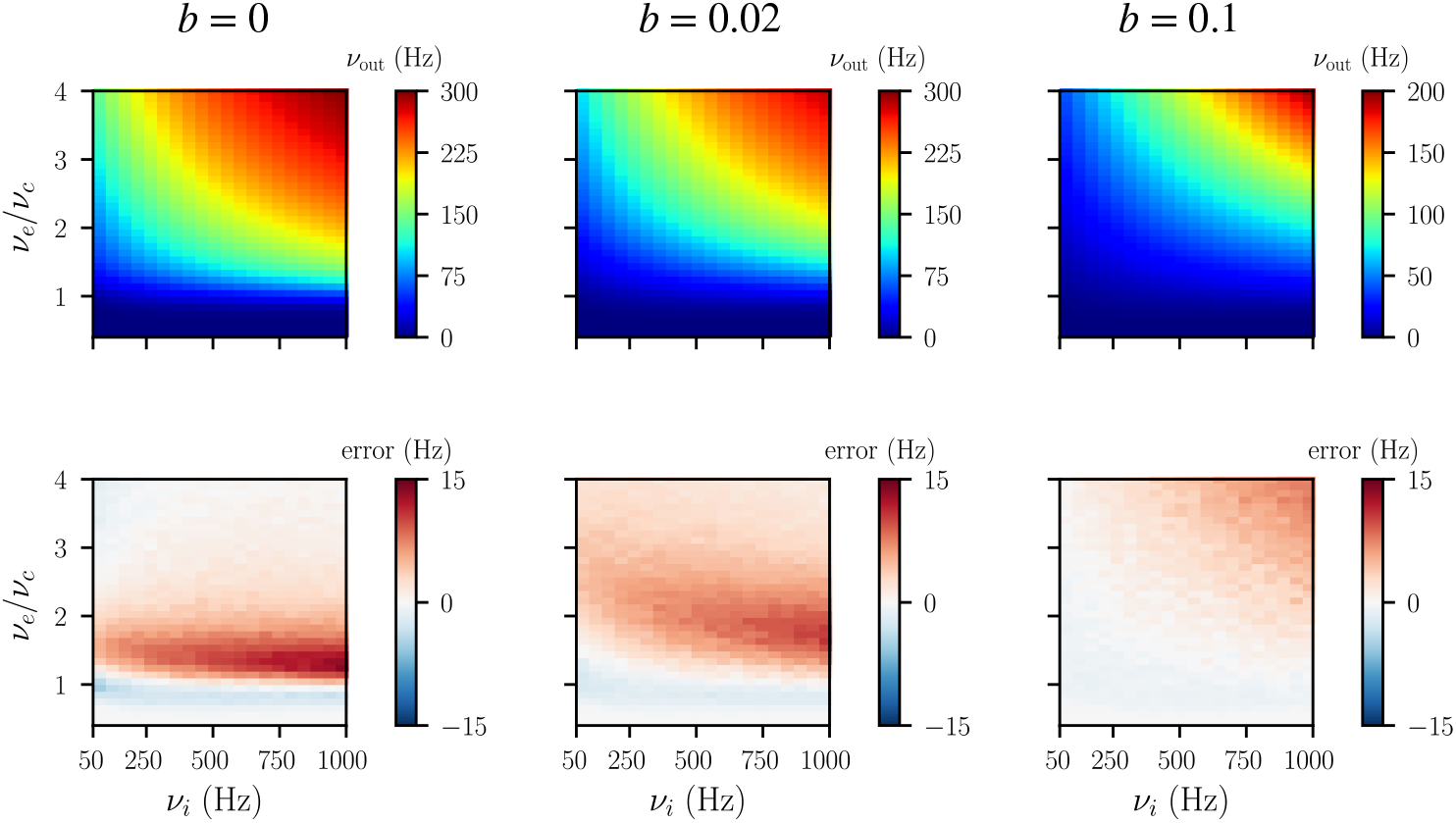
Accuracy of the dNMM in estimating steady-state firing rates. Three different spike-frequency adaptation parameters *b* = 0,0.02 and 0.01 *μ*A were adopted for testing. The upper row shows the population-average output firing rates, *v*_out_, given by different combinations of excitatory and inhibitory input rates, *v_e_* and *v_i_*. Note that the scale of the excitatory input was normalized by *v_c_*, derived in Eq. (13), in those plots. The bottom row shows the corresponding estimation error by comparing with those from MC simulations.

### Performance of simulating dynamic firing rate responses

In this section, we assessed the accuracy of Eq. (8) in estimating dynamic firing rates of a single neuronal population in response to time-varying stimuli. In particular, we shed light on the importance of 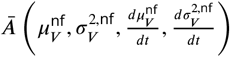 in Eq. (8) on estimating firing rates. Here, it was assumed that neurons only received nonstationary excitatory Poissonian spiking input, whose mean rate was described by an Ornstein-Uhlenbeck (OU) process,

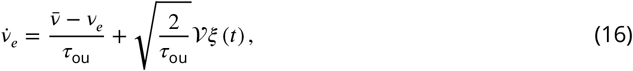

where *τ*_ou_ denoted the correlation time, 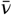 and 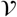 were the mean and standard deviation of the OU process, and *ξ*(*t*) was a unit Gaussian white noise process. Here, we set 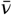 as 1.25*v_c_* and 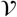 as 0.25*v_c_*, so that the input covered both the fluctuation- and mean-driven operating regimes. *τ*_ou_ was considered as the control parameter, which ranged from 10 to 100 ms. We adopted two performance measures, as in (***Ostojic et al., 2011***). One was the absolute difference between the estimated firing rates obtained from the dNMM and MC simulations, represented by root mean square (RMS) distance,

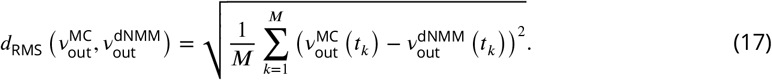

Here, 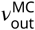 and 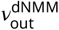 were respectively time series of the estimated firing rates from the MC simulations and dNMM, and each of them had *M* discrete elements over the calculating interval.

The other one was Pearson’s correlation coefficient:

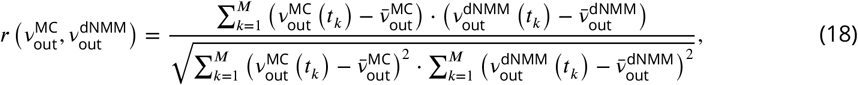

where 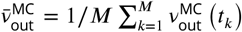 and 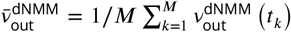 were time averages of discretely given firing rates.

We found thatthe dNMM with the complete rate function, i.e., Eq. (8), could accurately reproduce the time series of firing rates 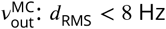 and *r* > 0.95 over the explored range of correlation times *τ*_ou_, regardless of spike-frequency adaptation strengths (see blue lines in ***Figure 4A***). However, when the term *Ā*(*) was ignored, the estimation accuracy became worse, especially for the cases where *τ*_ou_ = 10 ms (see red lines in ***Figure 4A***). Smaller *τ*_ou_ meant more rapidly changes in input rates. In other words, the term 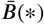 could not react quickly to rapidly changing inputs. As seen in ***Figure 4B***, especially for *b* = 0 *μ*A, the time series of the estimated firing rates from the incomplete rate function without *Ā*(*) had a phase lag as compared to that from MC simulations, while the complete rate function encompassing *Ā*(*) as well 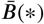 matched much better with MC simulations. Large spike-frequency adaptation (*b* = 0.1 *μ*A) decreased the sensitivity of neuronal populations reacting to stimuli, thus leading to increases in estimation accuracy of the incomplete rate function. We also observed that the estimation accuracy of the incomplete rate function became better for the case of *τ*_ou_ = 100 ms because of the more smoothed variation of the time-varying input (see ***Figure 4C***). But, it still could not capture rapid changes of output firing rates in this case.

**Figure 4.**
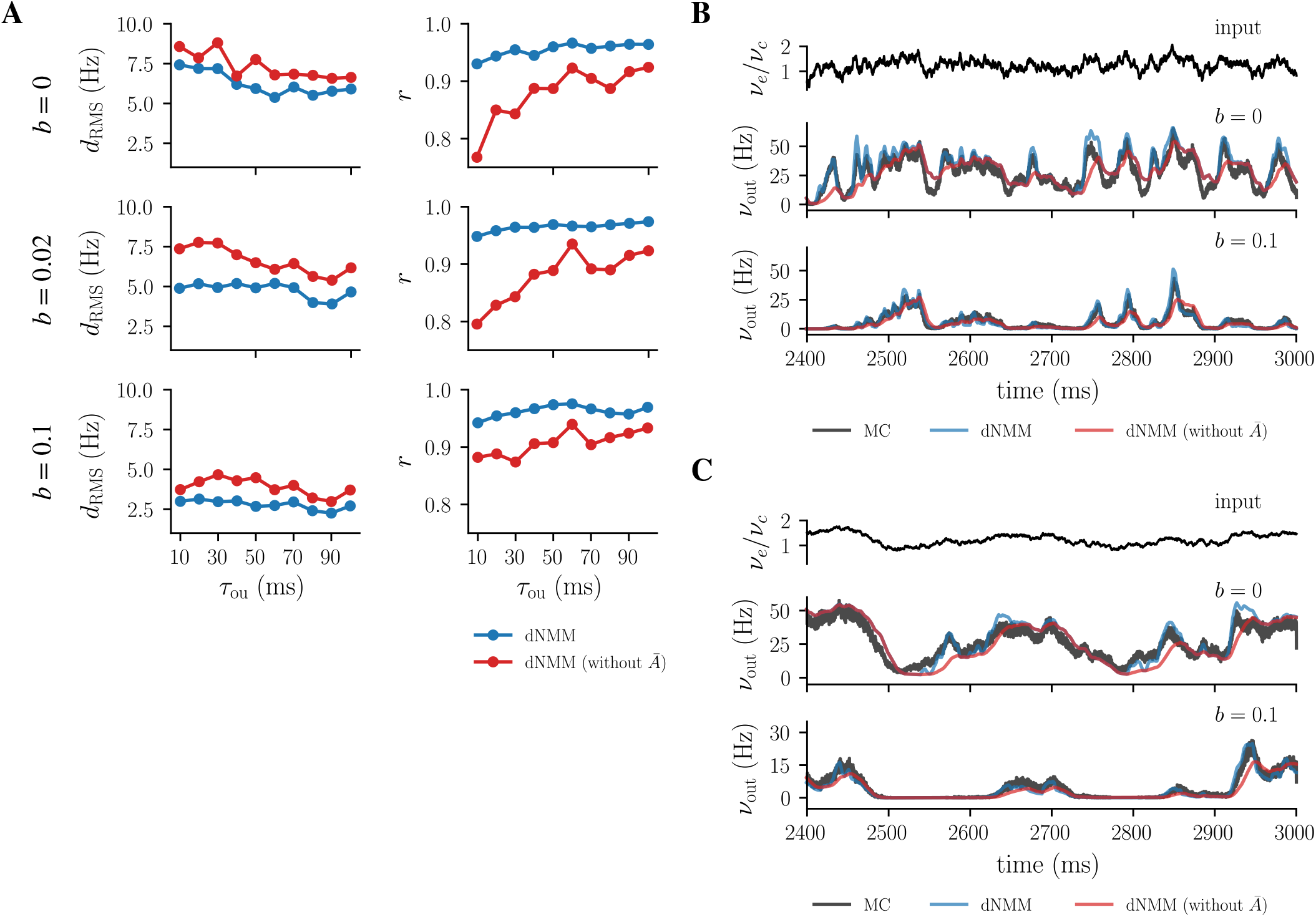
Accuracy of the dNMM in estimating dynamic firing rates in response to time-varying input represented by an OU process. (**A**) Root mean square distance, *d*_RMS_, and Pearson correlation coefficient, *r*, between the firing rate time series, 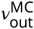 from MC simulations and 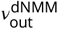 from the dNMM, for different spike-frequency adaptation strengths and different correlation times of the delivered OU input rate. The blue lines denote the performance using the complete rate estimate function, i.e., Eq. (8), while the red ones denote the performance using the incomplete rate estimate function where *Ā*(*) is ignored. (**B**) Examples of the time series of estimated firing rates. The upper row shows the delivered OU process with *τ*_ou_ = 10 ms. The middle one shows the firing rate time series from MC simulations (gray line), dNMM with a complete rate function (blue line) and an incomplete rate function (red line) under *b* = 0 *μ*A. The bottom one shows the time series under *b* = 0.1 *μ*A. (**C**) Similar to (**B**), except for *τ*_ou_ = 100 ms. OU process: Ornstein-Uhlenbeck process.

### Spike-frequency adaptation stablized asynchronous irregular activity of cortical networks

In this section, we moved on to the cortical network dynamics orchestrated by both the excitatory and inhibitory neuronal populations. Experiments showed that during the state of vigilance, intracellular recordings of cortical neurons in the awake animal revealed irregular discharges, and the neurons fire tonically at low firing rates (***Steriade et al., 2001; Destexhe et al., 2003***). In the viewpoint of network dynamics, such a network activity was referred to as the hallmark of the “asynchronous irregular” (AI) state, during which the population-average output firing rates sustainedly remained constant (***Brunel, 2000***). Typically, during this state, the inhibitory neurons had larger firing rates (around 30 Hz in average) compared to the excitatory neurons (around 10 Hz in average), which might result from the presence of spike-frequency adaptation in the excitatory neurons. This raised a question of whether the spike-frequency adaptation influenced the emergence of the AI states of cortical networks. Here, we attempted to study this question.

In this study, a local cortical networks was assumed to be characterized by a recurrent excitatory-inhibitory network composed of a pyramidal (PY) neuron population and an interneuron (IN) population, as in (***Brunel, 2000; Kumar et al., 2008; Augustin et al., 2013***). The details about the structure of this network is described in the section of Methods and Materials. We explored the network activity of such a recurrent cortical network as a function of two control parameters, the external excitatory input *v*_ext_ and the relative inhibitory strength *S*_inhi_ (i.e., Eq. (15)) by using the dNMM, and then detected the regime covered by the AI state in this parameter space. Results showed that the AI state emerged due to sufficiently strong *S*_inhi_, and the presence of spike-frequency adaptation (*b* > 0) allowed the AI state to cover the regime of lower *v*_ext_ and *S*_inhi_ (color shaded areas in ***Figure 5A***). Within the covered parameter space, the population firing rate of the PY population, *v*_out,PY_, for the AI state ranged from 8 to 25 Hz in the absence of spike-frequency adaptation, while it ranged from 5 to 22 Hz for *b_PY_* = 0.02 *μ*A and from 2 to 15 Hz for *b_PY_* = 0.1 *μ*A (characterized by colors in ***Figure 5A***). These magnitude levels of output firing rate roughly matched experimental observations of the AI network activity. Some examples of firing rate time series are shown in ***Figure 5B***(*b_PY_* = 0.02 *μ*A) and ***Figure 5C***(*b_PY_* = 0.1 *μ*A), showing that the dNMM gave a good estimation of AI cortical network activity comparable to MC simulations. Importantly, the presence of spike-frequency adaptation enlarged the regime covered by the AI state in the parameter space. This suggested that the spike-frequency adaptation in the excitatory neurons could stablize the AI network activity in the presence of fluctuating neuronal or synaptic properties.

**Figure 5.**
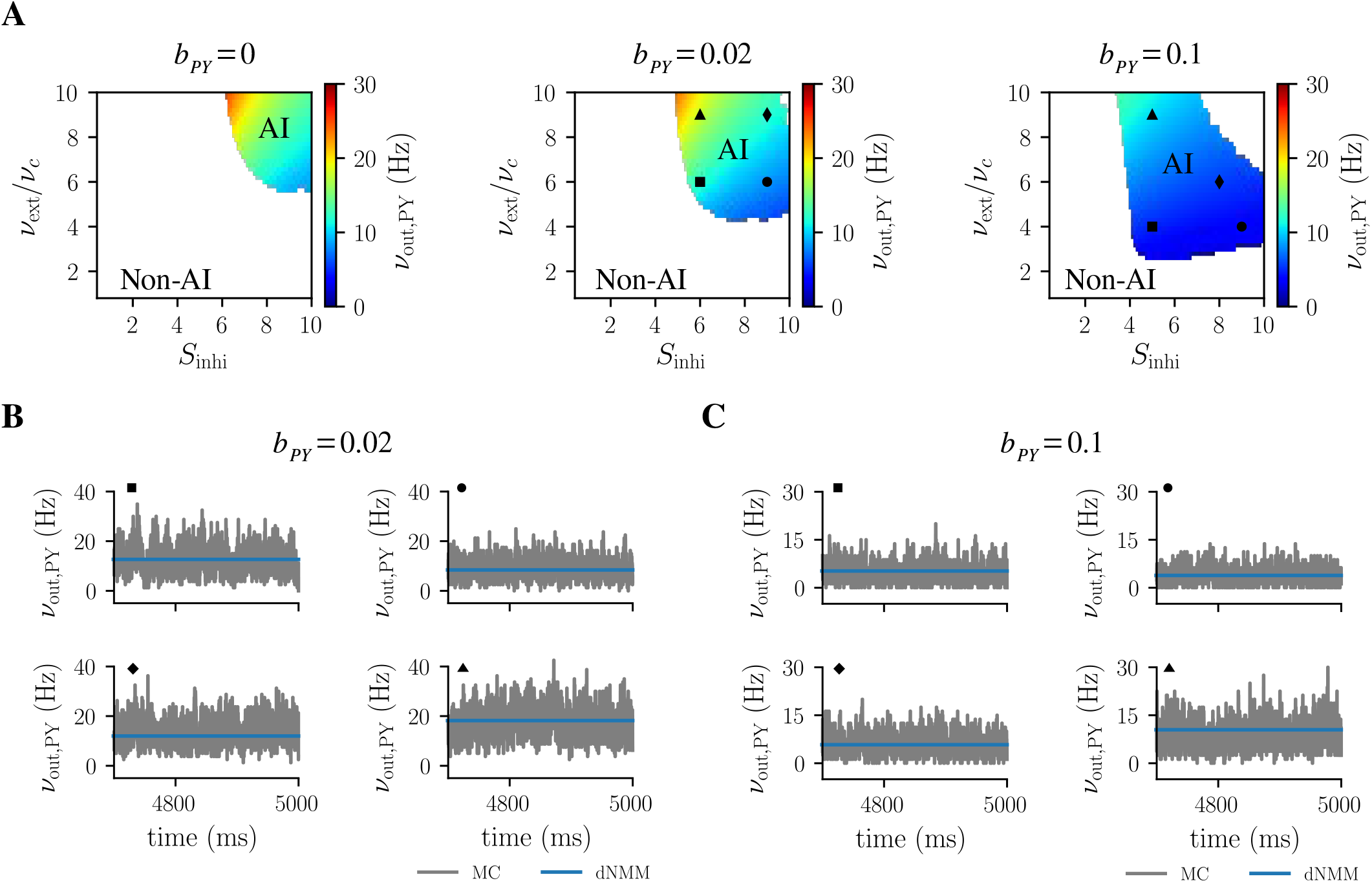
Characteristics of the AI state with respect to different spike-frequency adaptations. (**A**) Covered regimes of the AI state in a parameter space spanned by the external excitatory input *v*_ext_ (normalized by *v_c_*) and the relative inhibitory strength *S*_inhi_ for *b_PY_* = 0,0.02 and 0.1 *μ*A from left to right panel. Colored shaded areas: regimes covered by the AI state. Blank area: regimes covered by the non-AI state. *v*_out,PY_: the population firing rate of the PY population. (**B**) and (**C**) show four examples of firing rate time series estimated by the MC simulations and dNMM during AI states for *b_PY_* = 0.02 and 0.1*μ*A, respectively. AI stateL asynchronous irregular state.

## Discussions

In this study, we have developed a novel type of NMMs, termed dNMM, for simulating network dynamics of aEIF neuronal populations with biologically conductance-based synaptic interactions and adaptive behaviors of firing rate responses. Our study has demonstrated that, as compared to MC simulations, the dNMM gave accurate estimations of firing rate responses under both steady and time-varying input conditions. The dNMM could also correctly predict the activities of large scale neural networks consisting of multiple neuronal populations, such as the prediction of the regime having AI network activity of the cortical network shown in ***Figure 5***. In the future, like conventional NMMs, we will test the applicability of the dNMM to simulate other electrophysiological measurements of neural systems, such as the emergence of various rhythms in electroencephalography (***Ursino et al., 2010; Zavaglia et al., 2010; Cona et al., 2014***) or the changes of BOLD signals in functional magnetic resonance imaging (***Deco et al., 2013; Friston et al., 2017***).

Our dNMM tackled two major issues in current computational neuroscience community. First, the dNMM can be viewed as the next-generation of NMMs when one seeks new types of NMMs with the adaptive property (***Coombes and Byrne, 2019; Byrne et al., 2020***). Because of the presence of rigorously theoretic bottom-up frameworks based on the csPDM, dNMM overcomes drawbacks in the existing NMMs. For example, it retains all parameters of the underlying neuron model, such that one can explore which neuronal property predominates the superficial network dynamics, such as the spike-frequency adaptation that plays an important role in determining the emergence of AI network activity. A major discrepancy between the dNMM and other NMMs existing is the different rate functions used for estimating the population-average firing rates. Unlike the sigmoid rate function used in other NMMs, our rate function, i.e., Eq. (8), is based on the description of detailed spiking neurons (see Methods and Materials). As shown in ***Figure 2***, our rate function gives more accurate estimations of steady-state firing rates when compared to the sigmoid rate function. The dNMM, therefore unlike the convectional NMMs, is not just a phenomenological modeling approach.

The dNMM was tested by comparing its estimations of firing rate responses to that of the full spiking network model, i.e., the MC simulations, as shown in ***Figure 3, Figure 4*** and ***Figure 5***, and it gives accurate estimations. The rate function in the dNMM includes two terms: *Ā*(*) and 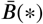. In physiological interpretation, the activity *Ā* arises as a result of threshold crossings by the mean and variance of membrane potentials caused by time-varying input, whereas the activity 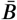 occurs due to threshold crossings by steady fluctuation- or mean-driven input. Similar forms of such a rate function can also be seen in previous firing rate models (***Chizhov et al., 2007; Buchin and Chizhov, 2010***), but they did not consider the threshold crossings by transient variances of membrane potentials caused by the changing input. We find that *Ā* helps to quickly react to the rapidly changing input, especially at the transition to the activated firing regimes from the non-firing regimes (as seen in ***Figure 4***).

It should also be noted that the estimation of *v*_out_ neglected the spike generation mechanism of aEIF spiking models. That was acceptable because the quantity of interest was the populationaverage firing rate determined by the number of the generated spikes of activated neurons, rather than their spiking times determined by the spiking generation mechanism. Furthermore, re-crossing of the spiking threshold voltage for activated neurons was actually involved by the expression of 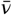 in the 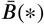.

The second issue tackled by dNMMs is the calculation inefficiency of PDMs (***Tranchina, 2009***). Due to the presence of only ordinary differential equations, the dNMM has excellent calculation efficiency on neural network modelings. In this study, we employed the trapezoidal method to solve all necessary ordinary differential equations and the midpoint method to calculate the population firing rates numerically. The function and constant that needed to be determined before simulation were the current-rate relationship of EIF neurons, *f*_EIF_(*I*), and the threshold voltage *V_th_*. It took around 12 seconds to simulate 10-seconds activity of the two-population cortical network with a time step size of 0.1 milliseconds (implemented in C++ programming with a CPU: i5-6500, 3.20GHz),

Our work is based on the csPDM (***Huang and Lin, 2018***), which we have augmented by explicitly including the dynamics of adaptation through aEIF neuron models. Due to its ability to reproduce a variety of firing patterns (***Naud et al., 2008; Touboul and Brette, 2008***), the use of the aEIF neuron model allowed us to simulate the activity of some brain regions where neurons revealed other special firing patterns, rather than just RS or FS pattern. The thalamus, for example, was in general the simulated target for studying slow-wave oscillations during sleep (***Destexhe, 2009; Crunelli et al., 2012; Cona et al., 2014***), where thalamic nuclei exhibited strong low-threshold spiking bursts (***Destexhe and Sejnowski, 2003***). In addition, following the original csPDM where the short-term plasticity of synaptic dynamics was encompassed, it is thus expected that, by just adding associated dynamic equations of the short-term plasticity, dNMM also can be used to simulate specific network dynamics, for examples, the persistent activity in working memory models (***Barak and Tsodyks, 2014***), that is significantly influenced by the short-term plasticity between synaptic interactions.

The premise of our dNMM is the Gaussian distribution of membrane potential variables, a solution of Fokker-Planck equation. This assumption can be seen in the prediction of the proportion of activated neurons (Eq. (9)) and of the effective current input to neurons (Eq. (10)). In turn, it also dominates the estimation of population firing rates. This is why the firing rate can be correctly estimated if such a premise is satisfied, even though the fluctuation of synaptic conductances cannot be well described by using diffusion approximations (see in ***Figure 1***). However, the satisfaction of this premise may depend on the choice of spiking neuron models. It is an adequate assumption for the integrate-and-fire neuron models, but does not guarantee its applicability to the integrate-and-fire-or-burst neuron models (***Smith et al., 2000***) or Izhikevich’s models (***Izhikevich, 2003***), which do not result in Fokker-Planck formalisms.

Using low-dimensional mean-f eld models to simulate large scale neural networks is a current trend in the computational neuroscience community (***Breakspear, 2017***). In terms of performance measures, *d*_RMS_ and *r*, the accuracy of the dNMM is comparable to other mean-field models, such as linear-nonlinear models (***Ostojic et al., 2011***), spectral models (***Augustin et al., 2017***) and master equation formalisms (***di Volo et al., 2019***), when, in general, *d*_RMS_ < 10 Hz and *r* > 0.95. However, synaptic interactions in the models by ***Ostojic et al. (2011)*** and ***Augustin et al. (2017)*** were current-based, which did not take the nonlinearity resulting from voltage-dependent synapses (***Paulus and Rothwell, 2016; Zerlaut and Destexhe, 2017***) into account. And the model by ***di Volo et al. (2019)*** only formulated the dynamics of firing rates, but did not include other state variables such as membrane potentials that might be relevant to electroencephalographical signals.

## Methods and Materials

In this section, the cortical excitatory-inhibitory network of aEIF neurons and details about aEIF neuron models were described, and the dynamic equations of the dNMM were derived step by step, starting from the descrption of the csPDM.

### The recurrent cortical network model

As proposed by ***Brunel (2000)***, here, a local cortical network was hypothesized to consist of both the excitatory PY and the inhibitory IN neuronal populations, in which the probability of connection between two neurons was 5%. The block diagram of this network circuit is shown in ***Figure 6***. We set there were totally *N* = 10000 neurons in this network. Eighty percent of them were considered to be excitatory, and the rest were inhibitory. So, each neuron received recurrent inputs from 400 excitatory PY neurons and 100 inhibitory IN neurons. In addition, each neuron also received excitatory inputs outside the network through 400 connections. We considered that the excitatory input was mediated through AMPA-receptors, and the inhibitory one was mediated through GABAa-receptors. The time scales of dynamics of both types of receptors were assumed to be short, at the level of milliseconds or tens of milliseconds (***Dayan and Abbott, 2001***). In this study, we set the strength of external excitatory inputs and the relative strength of recurrent inhibitory input as control parameters to explore the network activity. The delayed time for delivering spikes between neuronal populations was set as 2 ms.

**Figure 6.**
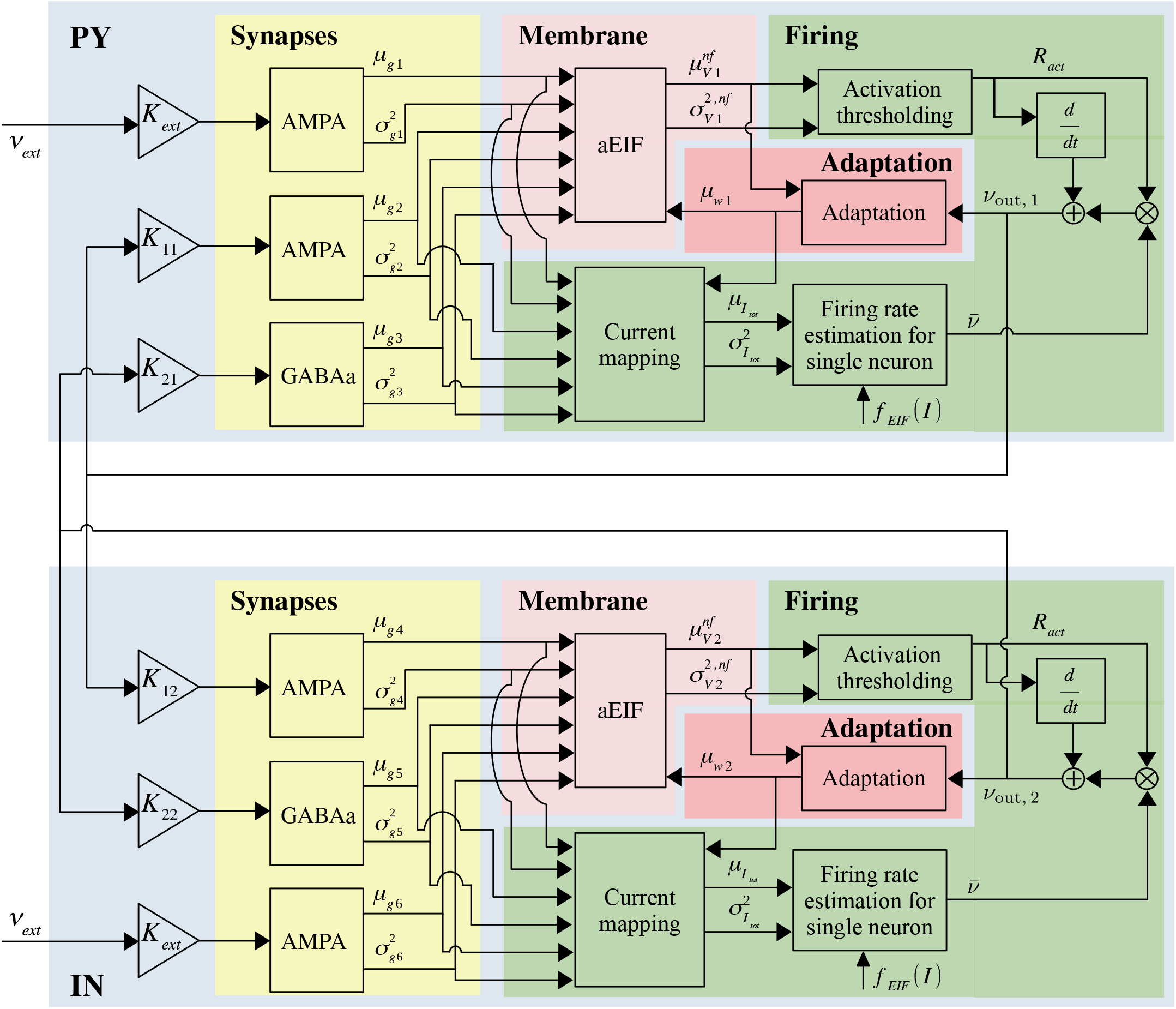
Block diagram of a local cortical network. The network consists of pyramidal (PY) and interneuron (IN) populations, both represented by aEIF models. Each neuron receives external excitatory input through AMPA-mediated synapses, and, meanwhile, receives excitatory and inhibitory recurrent connections from PY and IN populations, respectively, through AMPA- and GABAa-mediated synapses. *K_mn_* means the average number of connections from the *m* population to the *n* one, with *m, n* assuming 1 (PY) and 2 (IN). *K_ext_* means the number of connections from the external sources, and *v_ext_* is the mean rate of Poissonian input spike trains. In the dNMM the dynamics of each neuronal population is characterized by a serial combination of synaptic dynamics, describing time-varying means (denoted by *μ_gs_*, *s* ∈ {1,2,3,4,5,6}) and variances (denoted by 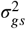, *s* ∈ {1,2,3,4,5,6}) of synaptic conductances, membrane dynamics, describing the time-varying means (denoted by 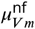) and variances (denoted by 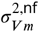) of membrane potentials, adaptation dynamics, describing the time-varying means (denoted by *μ_wm_*) of adaptation currents, and an estimator of the population-average output firing rate (denoted by *v_out, m_*).

### The spiking neuron model

Each neuron in the cortical network was modeled by using aEIF model (***Brette and Gerstner, 2005***), which combined the exponential integrate-and-fire (EIF) spiking neuron model (***Fourcaud-Trocmé et al., 2003***) with an adaptation current for capturing adaptive firing responses. The resulting dynamic equations were:

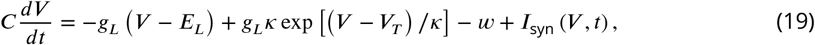

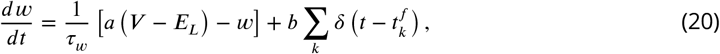

where *C* =1 *μ*F was the membrane capacitance, *g_L_* = 0.05 mS was the leak (resting) conductance, *E_L_* = −65 mV was the leak reversal potential, *κ* = 2.5 mV was the steepness of the exponential driving force to threshold, and *V_T_* = −50 mV was the spiking threshold voltage. Regarding the spike generation, when the membrane potential *V* reached the threshold, a spike was emitted, and *V* was then instantaneously reset to the resetting voltage *V_r_* = −65 mV and clamped for a refractory period of *τ_r_* = 2.5 ms. *w* was the adaptation current variable, with a time constant *τ_w_* = 600 ms. The strength of the adaptation current was controlled by two parameters: *a* = 0.005 mS for sub-threshold adaptation and *b* (in *μ*A) for spike-frequency adaptation. The value of *b* was set as zero for IN neurons that had no spike-frequency adaptation. In this study, the value of *b* for PY neurons was a control parameter to explore its associated effect on the types of network activity.

*I*_syn_ was the postsynaptic current, accounting for synaptic interactions induced by the presynaptic spikes. We modeled it as the sum of excitatory and inhibitory currents, given by:

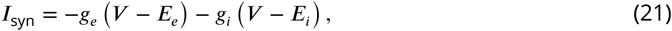

where *E_e_* =0 mV and *E_i_* = −80 mV were the excitatory and inhibitory reversal potentials, respectively. Here, we employed voltage-dependent conductance-based synaptic currents. *g_e_* and *g_i_* were synaptic conductance variables. We modeled their dynamics as first-order kinetics, in which the conductance was an exponentially decaying function that took instant jumps of amount Γ_*e*_ (Γ_*i*_) at each excitatory (inhibitory) presynaptic spike. Their dynamic equations could be written by (***Roth and van Rossum, 2009***):

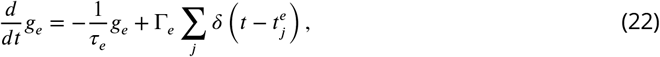

and

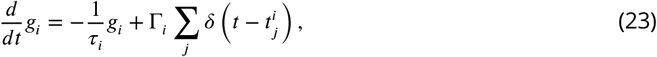

where *τ_e_* = 5 ms and *τ_i_* = 10 ms were synaptic time constants, 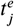 indicated the arrival time of the *j*-th presynaptic spike along the excitatory synaptic connections, and 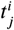 for inhibitory synaptic input. We considered that the arrival times, 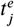 and 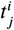, were governed by homogeneous or imhomogeneous stochastic Poisson processes. This assumption of Poissonian spike trains was in agreement with the findings of Poisson-like inter-spike interval distributions of cortical neurons under awake states (***Bédard et al., 2006; El Boustani et al., 2007***).

### The colored-noise population density method

Rather than the activities of individual neurons, dNMM offered a mean-field description of the collective activity of a sparsely connected random network of aEIF neurons as defined by Eqs. (19)–(23). Its derivation started from the so-called csPDM, in which the distribution of membrane potentials and the population firing rate could be calculated using the Fokker-Planck equation in the mean-field limit, i.e., *N* ≫ 1. A core assumption of the csPDM was that the synaptic dynamics operated under the diffusive limit, under which, in the presence of finite synaptic time constants, the stochastic fluctuating synaptic conductances could be approximated by nonstationary Gaussian colored noises. The corresponding dynamic equations for their means and variances were (equations (31) and (32) in ***Huang and Lin (2018)***):

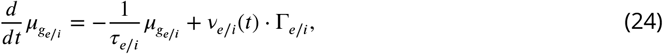

and

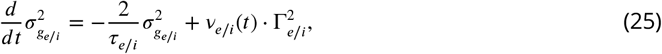

where *μ_g_*__* stood for the mean of the fluctuating conductance and 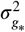 for the variance (***Figure 6*, Synapses**); *v_e_*(*t*) (*v_i_*(*t*)) denoted the time-varying mean rate of excitatory (inhibitory) presynaptic spike trains.

In order to follow the original idea of csPDM, where individual neurons were characterized by using the EIF spiking neuron model without adaptation, an adiabatic approximation was made on the adaptation current (***Augustin et al., 2013***). The adiabatic approximation was applicable if the time scale of the adaptation current was much slower than that of the membrane potential, i.e., *τ_w_* ≫ *τ_m_* = *C*/*g_L_*. Under this approximation, every neuron within the population had the same adaptation current, whose amplitude was approximated by its average over the population *μ_w_*(*t*). Accordingly, we yielded a new spiking neuron model equation when *μ_w_*(*t*) was substituted into Eq. (19), given by

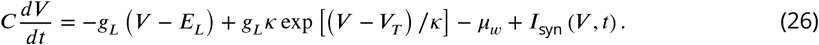

It turned out that Eq. (26) could be considered as a form of EIF neuron models if *μ_w_* was viewed as an additional current input. We therefore could directly apply the csPDM to Eq. (26). How to calculate *μ_w_* is described later in the Eq. (43). After applying the csPDM, we had the following partial differential equation to govern the temporal evolution of the density function (denoted by *ρ_V_*(*V, t*)) of the distribution of the membrane potential variables within the population:

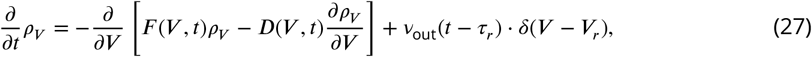

where

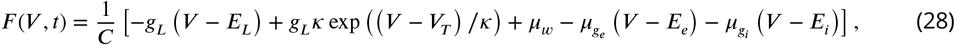

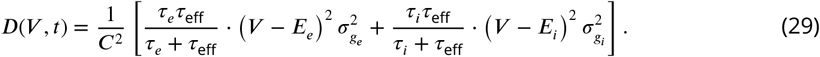

*v*_out_ denoted the population-average output firing rate of the neuronal population, *τ_r_* was the refractory period of individual neurons, and *τ*_eff_ was defined as the effective time constant of membrane potentials of conductance-based spiking neurons (***Gerstner et al., 2014***), given by

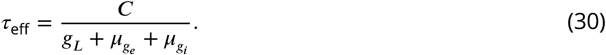

The details of derivations of these equations can be found in the paper of ***Huang and Lin (2018)***. In summary, the dynamic equations of the csPDM included a partial differential equation and a few ordinary differential equations, through which the population dynamics of a population of aEIF neurons receiving external excitatory and inhibitory inputs was described. It can be imagined, therefore, that, as compared to that for ordinary differential equations, most effort was made to solve the partial differential equation numerically when using the csPDM. This is one of the major motivations to develop dNMM.

### The density-based neural mass model

In this section, we moved on to the core work of this paper, i.e., the derivation of the dNMM based on the csPDM mentioned-above. First, we neglected the spike generation mechanism of individual neurons and, thus, defined a function 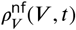, which described the distribution of membrane potential variables under this circumstance. Such a new density function evolved along with the time also according to a partial differential equation, similar to Eq. (27):

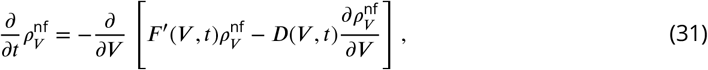

except for the removal of the re-entering source term, i.e., the last term in the Eq. (27). The boundary conditions of this equation were 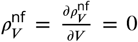 at both boundary sides of membrane potentials. Notice that in the Eq. (31), *F*(*V, t*) was replaced by

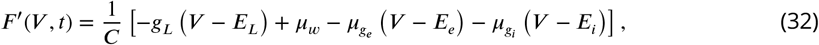

in which the exponential term was removed in order to avoid the mathematical divergence of the membrane potential when the spike generation mechanism was not taken into account. Finally, Eq. (31) was a typical form of nonlinear Fokker-Planck equations with state-dependent drift and diffusion coefficients. In other words, the solution 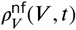 should be a nonstationary Gaussian function (***Risken, 1996***) which was characterized perfectly by its mean and variance.

According to the definition of the mean value 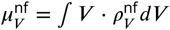, we yielded

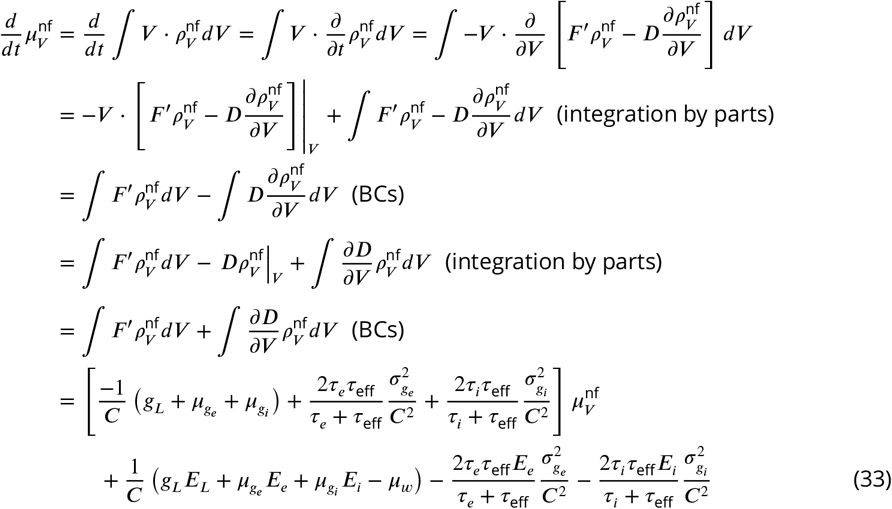

as the dynamic equation governing the time-dependent mean value 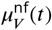. Similarly, we yield

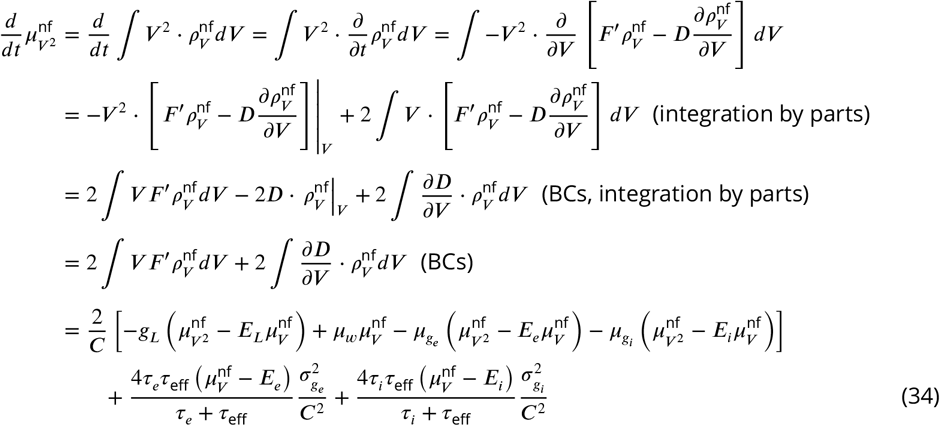

for governing the time-dependent mean value of the square of the membrane potential variable. Next, according to the identity 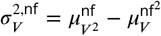, the dynamic equation for governing the variance 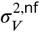 is given by

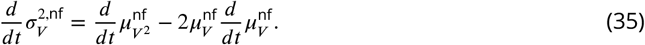

Replacing 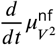 and 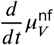 in the above equation with Eqs. (33) and (34), we then had

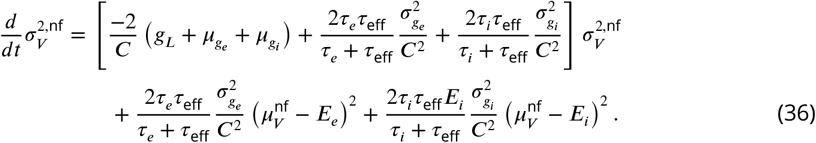

As a result, the partial differential equation, i.e., Eq. (31), was turned into two coupled ordinary differential equations, one for tracing the mean value and the other one for tracing the variance (***Figure 6*, Membrane**). The next step was to calculate the output firing rate of neurons (***Figure 6*, Firing**).

***Gerstner et al. (2014)*** def ned the population-average output firing rate *v*_out_ as

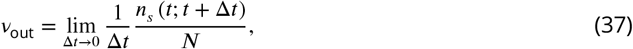

which was calculated through counting the number of spikes *n_s_*(*t*; *t* + Δ*t*) in a small time interval Δ*t* and dividing it by *N* and Δ*t*. Here, we assumed that the spikes within the interval Δ*t* came from two parts: (1) the initial firing of non-activated neurons from below the threshold, and (2) the re-enforced firing of activated neurons escaping from refractoriness if the input was still sufficiently large. Accordingly, *v*_out_ was calculated by

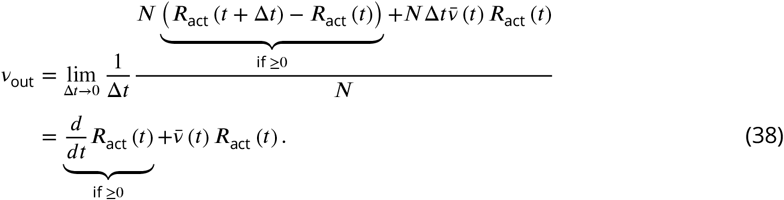

*R*_act_(*t*) meant the proportion of activated neurons whose membrane potentials exceeded a threshold voltage *V_th_* at time *t*, which was calculated as

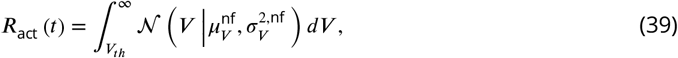

where 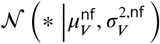 was a Gaussian density function with the corresponding mean 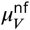 and variance 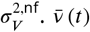 in Eq. (38) was assumed to be the mean firing rate of an activated neuron, and, as a result, 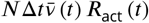 was the total number of spikes produced by activated neurons within the time interval Δ*t*. Here, 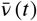 was estimated through a current-mapping algorithm. We assumed that the effective input current received by an activated neuron was characterized by a stochastic Gaussian process, whose mean was

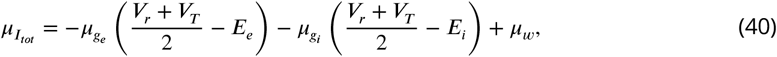

and whose variance was

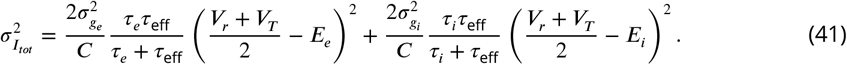

Assuming *f*_EIF_(*I*) was the current-rate relationship of the EIF neuron, then 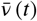 was given by

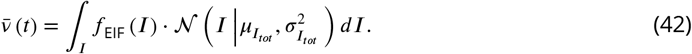

The calculations of *R*_act_ and 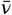 are illustrated in ***Figure 7***. What remained unknown was the dynamics of the mean adaptation current *μ_w_* (***Figure 6*, Adaptation**). According to the derivation in (***Augustin et al., 2013***), its dynamic equation was

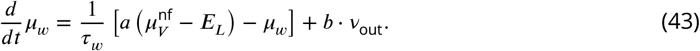

**Figure 7.**
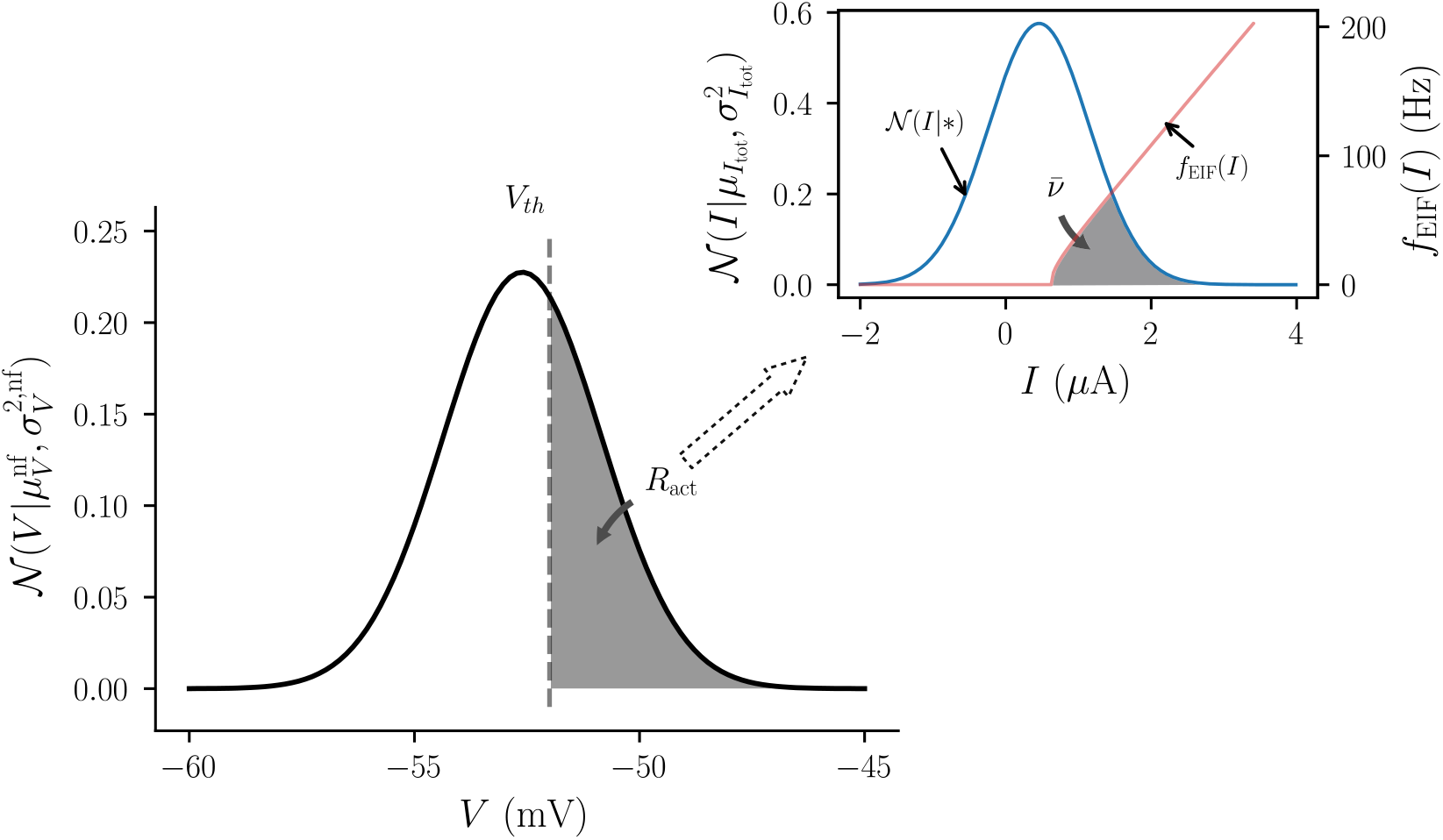
Illustrations of the proportion of activated neurons (*R*_act_) and the calculation of the average firing rate of activated neurons 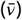.

We have stated all necessary dynamic equations of the dNMM, i.e., Eqs. (24), (25), (33), (36) and (43), and the estimate function of population-average output firing rate, i.e., Eq. (38), based on a single population of aEIF neurons. The idea of dNMM can be easily extended to random and sparsely connected neural networks consisting of multiple neuronal populations, like the cortical network mentioned above.

Notice that we did not directly consider the spike threshold voltage, *V_T_*, of the aEIF neuron as the threshold voltage for determining *R*_act_ because that the nonlinear exponential term and the spike generation mechanism were neglected. We hoped that these mismatches could be compensated by a careful choice of *V_th_*. So, in the present study, the value of *V_th_* was determined through optimally minimizing the error of estimating steady-state output firing rates given by a series of inputs. In our simulated cases, we set *V_th_* = −50.790 mV for the case of *b* = 0 *μ*A, *V_th_* = −50.907 mV for *b* = 0.02 *μ*A and *V_th_* = −51.133 mV for *b* = 0.1 *μ*A.

## Acknowledgments

This study has been supported by the Ministry of Science and Technology, Taiwan, Republic of China under grand no. 108-2321-B-006-024-MY2.

